# Long-term changes in populations of rainforest birds in the Australia Wet Tropics bioregion: a climate-driven biodiversity emergency

**DOI:** 10.1101/2021.07.23.453540

**Authors:** Stephen E. Williams, Alejandro de la Fuente

## Abstract

Many authors have suggested that the vulnerability of montane biodiversity to climate change worldwide is significantly higher than in most other ecosystems. Despite the extensive variety of studies predicting severe impacts of climate change globally, few studies have empirically validated the predicted changes in distribution and population density. Here, we used 17 years of bird monitoring across latitudinal/elevational gradients in the rainforest of the Australian Wet Tropics World Heritage Area to assess changes in local abundance and distribution. We used relative abundance in 1977 surveys across 114 sites ranging from 0-1500m above sea level and utilised a trend analysis approach (TRIM) to investigate elevational shifts in abundance of 42 species between 2000 – 2016. The local abundance of most mid and high elevation species has declined at the lower edges of their distribution by >40% while lowland species increased by up to 190% into higher elevation areas. Upland-specialised species and regional endemics have undergone dramatic population declines of almost 50%. The “Outstanding Universal Value” of the Australian Wet Tropics World Heritage Area, one of the most irreplaceable biodiversity hotspots on Earth, is rapidly degrading. These observed impacts are likely to be similar in many tropical montane ecosystems globally.

## Introduction

There is widespread recognition that climate change is rapidly becoming the most significant threat to global biodiversity and natural ecosystems [1, 2]. At a global scale, estimates of total species extinctions projected over the remainder of the century vary considerably between ecosystems, taxa and methods of analysis [3]. However, in all studies, the level of predicted impacts is disturbing with potential losses of between 15-35% of all species [3, 4]. The projected impact of climate change is expected to be especially severe in mountain ecosystems with up to 84% of mountain species globally facing a high extinction risk [5–7]. Mountain regions provide a host of critical ecosystem services [8], support a disproportionate amount of the world’s biological diversity and, harbour many phylogenetically unique species [5, 9–12]. Mountain regions contain roughly 87% of the world’s vertebrate biodiversity [13], 54% of which is completely restricted to mountain ecosystems [13]. The global significance of montane ecosystems is heightened in the tropics [14, 15]; approximately 50% of the world’s species of plants and vertebrates are believed to be endemic to 34 identified global biodiversity hotspots [16], 85% of which include large areas of tropical forest or montane cloud forest [17]. Tropical species are considered to be particularly sensitive to climate change [18–20] and the warming rates are relatively high in tropical mountains [18]. Consequently, tropical montane biodiversity is not only globally important but particularly threatened [7, 21]. Despite the global significance and high vulnerability of tropical ecosystems, there have been few studies demonstrating observed impacts of climate change in the tropics [22, 23]. The paucity of tropical studies makes it difficult to measure and predict the impacts of climate change relative to other drivers like habitat loss [24], especially given that most studies are short-term or lacking abundance data [25]. There is a need for increasing monitoring and improving understanding of the impacts of climate change in tropical montane ecosystems [12].

On mountains, biotic communities and abiotic conditions change abruptly over short distances, with greater elevational than lateral turnover in species composition [26]. Across all elevations, assemblages on mountains with high rates of past temperature change exhibit more rapid diversification, highlighting the importance of climatic fluctuations in driving the evolutionary dynamics of mountain biodiversity [12]. Globally, increasing evidence indicates that species are responding to climate change by shifting their geographical distributions [27]. These shifts often follow warming temperatures poleward and upslope [6, 10]. Montane species are of particular concern in this respect, as they are expected to experience reduced distribution area, increased population fragmentation, and increased risk of extinction with upslope movement into ever-smaller area [28, 29]. The high degree of specialization to narrow temperature ranges that montane species typically exhibit has raised concern over their future under climate change [10, 29, 30]. It is widely expected that montane species will experience further upslope shifts in the future and, in the absence of broad latitudinal shifts due to the geographical features of montane ecosystems, such movements will leave species with less habitable area as they approach mountain peaks [29, 31]. Left with nowhere else to go, montane species are predicted to become increasingly susceptible to the stochastic extinctions or declining populations [32]. This so-called “escalator to extinction” [33] has been predicted, and now observed, in a number of places and taxa around the world [28, 34–36].

The rainforests in the Australian Wet Tropics bioregion in north-east Queensland are globally significant for high biodiversity value based on high endemism, evolutionary significance and phylogenetic distinctiveness [37–39]. These biodiversity values resulted in the region being listed as a World Heritage Area in 1988 and being described as the sixth most irreplaceable protected area globally [40] and the second most irreplaceable World Heritage Area [41]. The high endemism and relictual nature of the biodiversity within the region is largely attributed to the influence of historical fluctuations in rainforest area over the Quaternary and the restriction of rainforest to cool, moist, upland refugia [37]. This biogeographic history imposed a non-random extinction filter across the region resulting in most of the regionally-endemic species being cool-adapted upland species [42, 43]. It is this biogeographic history, with the resulting concentration of endemic species in the cool uplands, that has made the biodiversity of the region so unique but highly vulnerable to a warming climate. Predictions about the future of this biodiversity under anthropogenic climate change are grim, particularly for the upland regionally-endemic species and habitat types [5, 44–46]. In 2003, species distribution modelling of the endemic vertebrates suggested the potential for catastrophic impacts over the coming century with more than 50% of these species predicted to go extinct due to a complete loss of suitable climatic space [5]. These predictions drove a greater research effort in the region in the interim years and there have been extensive region-wide surveys of many vertebrate and invertebrate taxa [47–50].

Are the declines in species ranges predicted in 2003 concordant with observed spatial trends in species abundance patterns over subsequent years? Unfortunately, the answer is yes. We present quantitative evidence, based on the long-term monitoring of vertebrates across the entire bioregion, for significant declines and shifts in the spatial distribution of populations. Field monitoring clearly demonstrates that the previously projected impacts are clearly concordant with observed shifts in species abundance. Previous analyses of the climate change impact in the region have relied on either modelled distribution changes using various IPCC emission scenarios or coarse comparisons of changes pre-2008 compared to post-2008. Here, we examine in high spatial, temporal and taxonomic detail the observed changes in the rainforest bird assemblages of the region between 2000-2016, based on the regional-scale standardised surveys from the Williams Wet Tropics monitoring program (updated from Williams, VanDerWal (47)). We use this long-term dataset to test for bird species that have undergone significant changes in local abundance and/or elevational. We predicted that bird assemblages should systematically shift upwards in elevation and that the local abundance of individual species would decline on the lower (warmer) edge of their distribution and increase at the higher (cooler) edge of their distribution [31]. We tested for trends across time in local abundance (site/elevation specific) and assemblage shifts across elevation and used an area-weighted trend to examine trends in total population size. These impacts are likely to be representative of impacts in many mountain ecosystems across the globe [7].

## Results

Overall, across all 42 species over the 17 years, there has been a significant decline in local abundance of rainforest birds of approximately 12 ± 1.4% (~ −0.2% per year) (Figure 1a, Table 1). However, this overall trend masks complex, and often contrasting trends, within different ecological subsets of the rainforest bird assemblage (Table 1). Habitat generalists increased by more than 50% from 2000 to 2011 and then steadily declined until 2016 (13 species, 3.3%/year, overall trend 9 ± 4.1 %, Figure 1b, Table 1). Local abundances of rainforest specialist species have declined on average by approximately 20% (29 species, −1.7%/year, overall trend −20 ± 1.3 %, Figure 1C, Table 1). Regionally endemic species, a subset of habitat specialists, showed the strongest decline with a loss of ~34 ±1.7% in local abundance (10 species, −2.4%/year, Figure 1d, Table 1). Population trends of habitat generalists and specialists were significantly different (trend difference 0.05 ± 0.002, p<0.05). However, these average multi-species trends in ecological groupings also mask variable trends for individual species (Figure S1) and assemblages in different elevational bands (Table S3). Species-specific trends in local abundance and total population size (local abundance trends weighted by area within each elevational band) are presented in the Appendix (Figure S1) (Temporary link to interactive Appendix - https://alejandrodelafuente.shinyapps.io/BirdsPopTrendAWT/?_ga=2.148260535.1024527938.1618546081-1577712465.1581926346.

**Figure 1.**
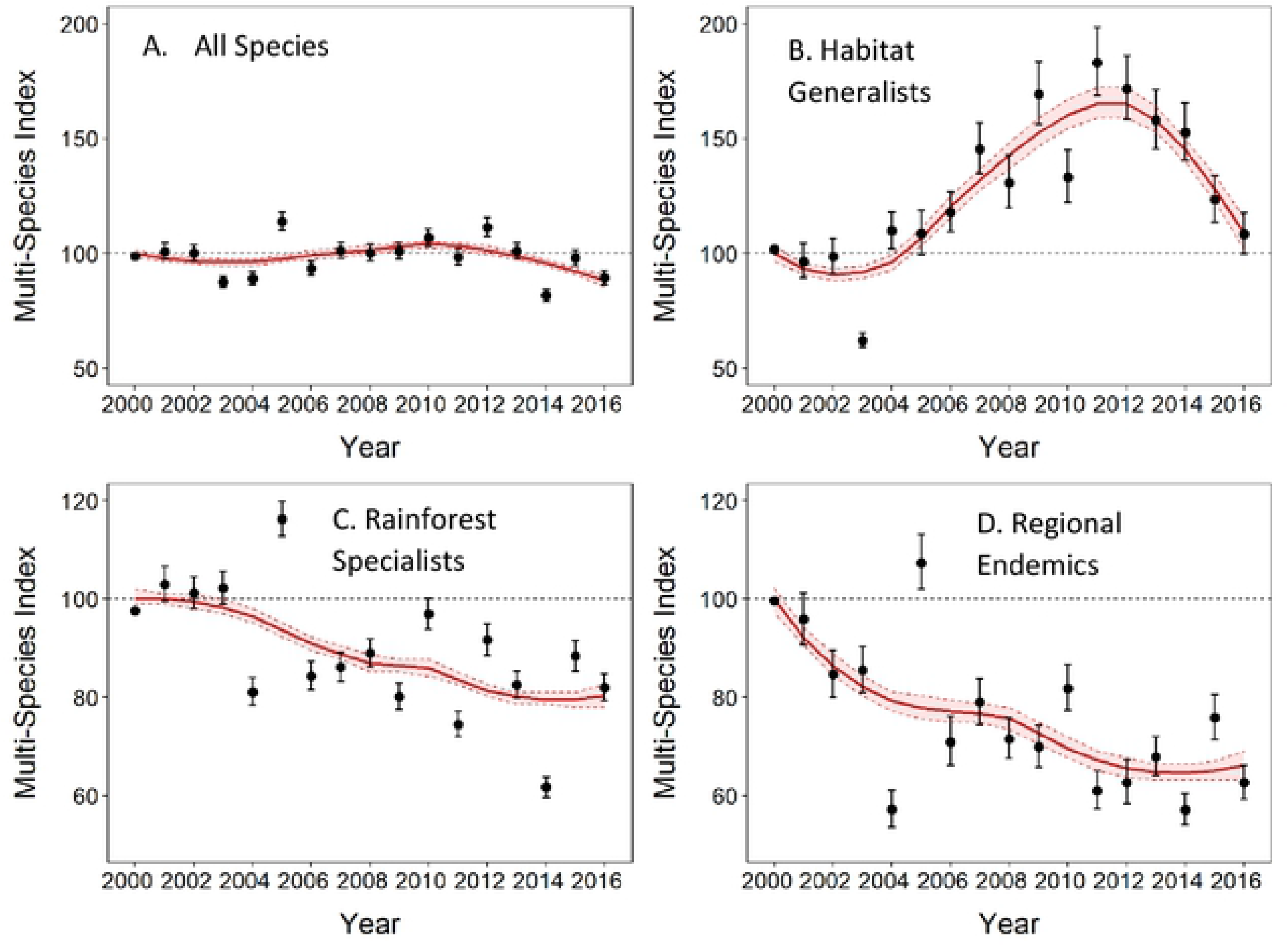
Trends in local abundance from 2000 to 2016 based on the multi-species indicators for a) all species, b) habitat generalists, c) regional endemic species and d) rainforest specialists. Values are the multi-species index with SE (error bars) with the smoothed trend line with 95% CI (shaded area).

**Table 1.** Multi-species indicator trends between 2000 – 2016 (overall) and separate trends pre-2009 and post-2008. Population trends were classified following Pannekoek and van Strien (51) into one of the following categories depending on the overall multiplicative slope and its 95% confidence interval. Strong increase/Steep decline: increase/decline significantly more than 5% per year. Moderate increase/decline: significant increase/decline, but no more than 5% per year. Stable: no significant increase or decline, and it is certain than trends are less than 5% per year. Uncertain: no significant increase or decline, but it is not certain if trends are less than 5% per year.

|  | Overall |  |  | 2000 - 2008 |  |  | 2009 - 2016 |  |  |
| --- | --- | --- | --- | --- | --- | --- | --- | --- | --- |
| *Indicator* | *Slope* | *s.d.* | *Trend classification* | *Slope* | *s.d.* | *Trend classification* | *Slope* | *s.d.* | *Trend classification* |
| All species | 0.998 | 0.001 | Moderate decline | 1.003 | 0.002 | Stable | 0.982 | 0.002 | Moderate decline |
| Endemic species | 0.976 | 0.001 | Moderate decline | 0.967 | 0.003 | Moderate decline | 0.985 | 0.004 | Moderate decline |
| Lowland species | 1.03 | 0.002 | Moderate increase | 1.074 | 0.005 | Strong increase | 1.008 | 0.005 | Stable |
| Midland species | 0.986 | 0.001 | Moderate decline | 0.992 | 0.003 | Moderate decline | 0.98 | 0.003 | Moderate decline |
| Upland species | 0.971 | 0.001 | Moderate decline | 0.952 | 0.003 | Moderate decline | 0.972 | 0.003 | Moderate decline |
| Habitat generalists | 1.033 | 0.002 | Moderate increase | 1.054 | 0.005 | Moderate increase | 0.974 | 0.005 | Moderate decline |
| Rainforest specialists | 0.983 | 0.001 | Moderate decline | 0.981 | 0.002 | Moderate decline | 0.986 | 0.002 | Moderate decline |

Lowland specialist species have undergone a strong and significant increase of 72 ± 7% (6 species, 3%/year, Figure 2a). Mid-elevation specialists declined by 21 ± 1.4% (16 species, - 1.9%/year, Figure 2b) and upland specialists have undergone declines of 44 ± 4% (13 species, −2.9%/year, Figure 2c). Population trends for lowland specialists were significantly different from the trends in both midland (trend difference 0.043 ± 0.002, p<0.05) and upland specialists (trend difference 0.059 ± 0.002, p<0.05). Additionally, upland specialists have declined significantly more than midland specialists (trend difference 0.015 ± 0.002, p<0.05), suggesting that the pattern of decline increases with increasing elevation.

**Figure 2.**
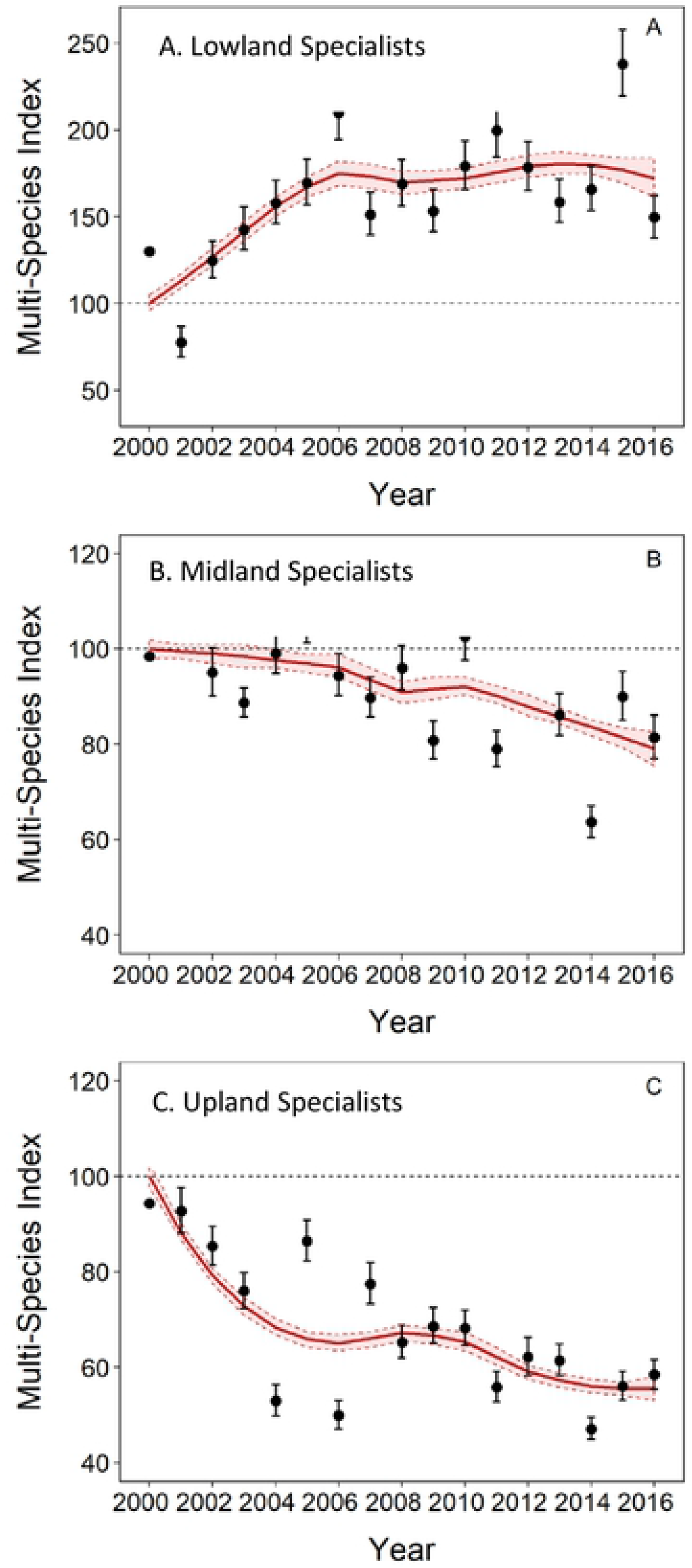
Trends in local abundance from 2000 to 2016 based on multispecies indicators for A) lowland specialists, B) midland specialists and C) uplands specialists. Values represent multi-species index with SE (error bars). The overall smoothed trend is represented by the line with shaded area showing the 95% CI for the smoothed trend.

Shifts in patterns of local abundance are species-specific and highly variable, often involving complex spatio-temporal and non-monotonic trends (for example, Brown Gerygone, Figure S1, Figure S1.1). However, the overall multi-species trends that summarise the observed shifts in bird assemblages along the elevational gradient are what we would predict under a warming climate: there have been significant upslope shifts in bird abundance patterns across the 17 years of this study. We demonstrated this by examining the trends in local abundance for each group of elevational specialists in their original preferred elevation (baseline averaged abundance at each elevation across 1996-2003) and in the adjacent elevational bands over time. We predicted, for example, that lowland species should increase in the midlands, and upland species should decline at lower elevations. The observed spatial shifts in elevational distribution and abundance are in accord with the earlier predictions based on distribution changes. Lowland specialist species have moderately increased in local abundance in the lowlands (+17%). On the other hand, lowland specialists’ local abundance has dramatically increased into the midlands (~190% increase) (Figure 3). Midland specialist species have declined in the lowlands by approximately 42%, by 22% in the midlands and are currently stable in the uplands (Figure 3). Upland specialist species have declined everywhere, with a catastrophic 49% decline in the lower-elevation midlands and a 33% decline in the uplands (Figure 3). Detailed trends for individual species at each elevation category are in Appendix (Table S3).

**Figure 3.**
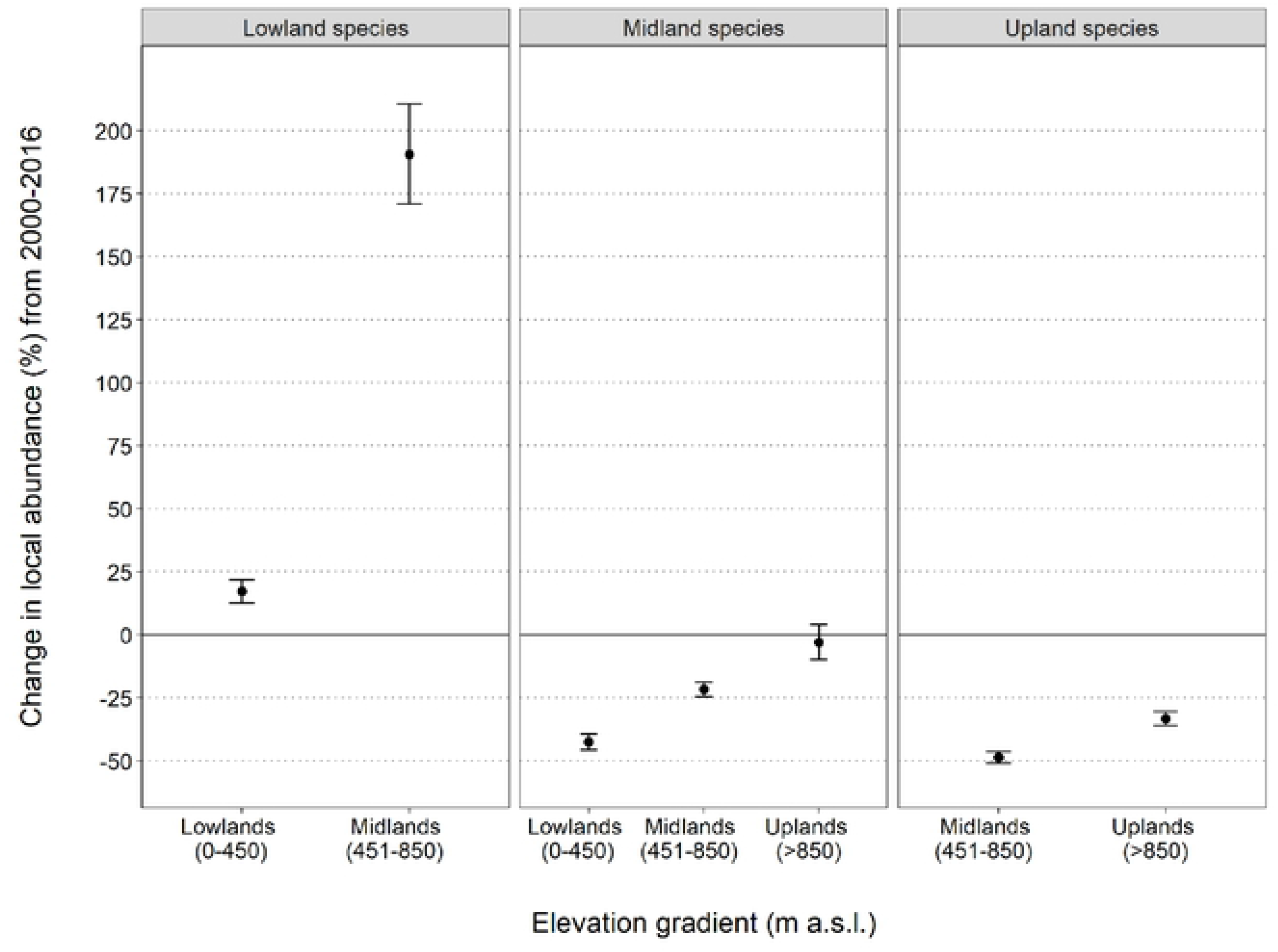
Elevatioπal shifts in local abundance patterns in bird assemblages between 2000 to 2016 across the elevatioπal gradient. Baseline population set to 0 with values above the baseline representing an increase in local population abundance and values below the baseline representing a decline. Values represent the change in local abundance over the full time period based on the mean multiplicative slope of the multi-species trend with its SE (error bars). See the full abundance trends over time for each combination of elevatioπal band and species group in Appendix (Figure S4, Table S4).

## Discussion

There is a temptation for policy makers and environmental managers to consider that biodiversity within a protected area is safe, this is a dangerous, and in this case incorrect, assumption. The montane rainforest birds of the Australia Wet Tropics World Heritage Area are in danger of extinction due to the increasingly severe impacts of a changing climate. In 2003, Williams et al. [52] predicted catastrophic extinction levels within the regionally endemic species of the Australian Wet Tropics World Heritage Area by the end of the 21^st^ century. These early predictions were based on bioclimatic envelope models of the changes in species distributions with increasing temperature. However, the reliability of predictions using this simple approach is debateable, with a likelihood of overestimating the severity of impacts [53]. Therefore, a more intensive monitoring effort was conducted throughout the region, covering ~94% of the environmental space present and providing standardised estimates of local abundance for many species.

An earlier analysis using these local abundance measures of rainforest birds across elevation [45] predicted that 74% of rainforest bird species would become threatened by the end of the 21^st^ century. demonstrated a tight relationship between elevation and the assemblage structure of birds based on empirical, site-based relative abundance across the elevational gradient. Based on this relationship, they predicted that bird assemblages would systematically move upwards in elevation as the climate warmed. However, despite the development of more sophisticated modelling approaches and the availability of more field data, the qualitative outcomes of predictions remained similar to the 2003 predictions. Here, we demonstrate that these predicted trends, whether based on species distribution models or empirical abundance, were concordant with expectations. Our results here demonstrate that the rainforest bird assemblage in the Australian Wet Tropics bioregion is clearly, and consistently, moving uphill in a classic demonstration of the “escalator to extinction” [33].

Overall, we are on track for severe impacts on rainforest birds, particularly the upland specialists. Many of these species are endemic to the region and/or include species that are evolutionary distinct and/or ecologically specialised [38]. Upland bird populations have declined since 2000 by 33% within their core range and the more marginal populations in the midlands have crashed by almost 50% (Figure 3). Midland species, although stable in the cooler uplands at this stage, have declined by >20% within their core area and >40% in the hotter lowlands. Lowland species are mostly increasing in abundance thus far with increases in abundance in the lowlands and strong increases in range and abundance into the midlands (Figure 3, Table 1).

The pattern in temporal change in abundance of rainforest birds presented in this study is consistent with observed species elevational shift worldwide [27, 34–36, 54–56]. Globally, evidence suggests a strong uphill elevational shift across different taxa, indicating that this trend could be a generality. However, some studies have pointed out some idiosyncratic results found in species-specific trends within different communities [27, 54]. According to this, our results showed that, even though the uphill distribution shift is marked across most species, some birds showed stable trends, while others showed pattern opposite than expected (e.g., *Acanthiza katherina*, figure S1). Those individual exceptions to the overall pattern highlighted potential interspecific differences in dispersal capacity, resilience, and adaptability between bird species in the Wet Tropics, whose impact in the overall community “re-shuffling” at the ecosystem level should be further studied. Overall, this study supports growing evidence tof the rapid impact that climate change is having on tropical ecosystems [28] and provides the first evidence of a climate-induced elevational shift in tropical rainforest of Australia.

So, what is driving these shifting abundance patterns? Is it the direct impacts of temperature on the birds or is it an indirect effect via food resources or other biotic interactions? There is little confidence in our ability to predict the potential impacts of the complex changes in biotic interactions due to these marked changes in abundance and geographic shifts in distributions. Upland and midland assemblages are likely coming under increased pressure due to changing biotic interactions and community structure associated with the increasing abundance of lower elevation species as they shift higher up the mountain. The changes shown in this study suggest that, thus far, the impacts on species have been largely direct, or at least directly associated with an important resource, rather than changes in competitive interactions between bird species. We argue this on the basis that upland bird species have declined in their core range (>850m) despite there being no significant increase in midland birds in the uplands thus far. Midland birds have only declined by about 20% in the midlands despite the influx of lowland species. Lowland species have increased (albeit with more recent trends of decline) and there has been no noticeable change in species composition or new invading species in the lowlands to date. Thus far, there is no evidence supporting “lowland attrition” of species in this system [31]; however, this is not entirely surprising given that we have little knowledge of the upper thermal limits of the lowland species because they already occur in the hottest part of the region Shoo, Anderson (57) suggested that the elevational pattern of abundance for the Lewin’s Honeyeater was directly influenced by temperature and not competitive interactions. This conclusion was based on a demonstration that the elevational abundance profile of the species varied as predicted by temperature in the populations on mountains on Cape York, a hotter montane system, within a very different bird assemblage to that in the Wet Tropics region[57]. However, this does not account for other biotic interactions that might also be changing such as diseases, or other non-bird competitors such as ectotherms (invertebrates mostly) that might be increasing in uplands.

There are many indirect mechanisms that could potentially exacerbate the impact that climate change will ultimately have on bird species. Increased pressure from parasites with increasing temperature [58], decreased reproductive success during prolonged dry seasons [59, 60], increased habitat and population fragmentation [61–63] and increased environmental marginality [64]. In this region, Williams, Shoo (50) hypothesised that dry season bottlenecks and changes in net primary productivity could have a strong influence on bird populations. Net primary productivity is limited by both temperature and water availability and could possibly increase in the uplands with a warming climate, potentially alleviating some of the more direct impacts of high temperatures. However, the strong declines in the abundance of upland bird species shown here suggests that any positive influence of increased net primary productivity has been swamped by the negative impacts of increasing temperature. There is existing evidence highlighting the importance of changing rainfall patterns, especially harsher dry-seasons [59, 65] and decreasing water input from cloud interception [66]. It seems most likely that the driving factor behind many of the changes demonstrated in this study are the increasing frequency and intensity of heat waves [67]. We need to increase our understanding of the impacts of extreme climatic events so we can make more robust predictions than those that rely only on changes in average conditions.

There have been clear impacts on biodiversity in almost every ecosystem and taxa across the globe due to anthropogenic climate change [2]; and now the world urgently needs to reduce emissions and adapt as much as possible to minimise future impacts. Our efforts need to be firmly focussed on the difficult question – what can we do? Managing habitats at the landscape scale via habitat restoration, threat abatement and enhancing dispersal pathways represents one avenue for local adaptation efforts to increase the resilience of biodiversity (Shoo et al. 2013, [68]). The maintenance and restoration of movement pathways and corridors to facilitate species movement into refugial areas will be vitally important [69], however, our results here suggest that facilitating movement also warrants caution. While it is imperative that many species can move into cool refugia [70], our data demonstrating the observed movement of lowland, generalist species into the upland refugia, represents a potential threat to the upland endemics via increased competition with more generalist species.

It is clear that montane systems are of paramount importance due to their high biodiversity value, many specialists and endemics and their role as the best-available cool refugia [70] and that these biodiversity hotspots are particularly threatened by climate. It is particularly disconcerting that, even in a fully protected and well-managed system such as the Australian Wet Tropics World Heritage Area, observed impacts are significant and accelerating. Most other tropical, montane biodiversity hotspots across the globe also face additional threats such as ongoing habitat degradation, poaching and urban encroachment.

## Conclusions

Upland bird species, of great conservation importance, are declining in abundance and contracting their range to higher elevations in the montane rainforests in the Australian Wet Tropics World Heritage Area in a classic example of the “escalator to extinction” [33]. These species are suffering the combined, and increasing, threat of reduced distribution area, reduced local population density and more fragmented and isolated populations, potentially causing a loss of genetic diversity in many species; factors that increase their vulnerability to extinction

[69]. Low-elevation rainforest species that are often more generalist, geographically widespread and locally common are increasing in abundance and range size, potentially resulting in yet another negative pressure on the upland specialists and an overall homogenisation of the rainforest avifauna [71]. These changes are likely to be indicative of impacts across all montane ecosystems globally, especially in important biodiversity hotspots associated with tropical mountain ecosystems. The important next step is to determine how we can slow, prevent, or adapt to these impacts to prevent the loss of the unique biodiversity of these regions around the world.

## Methods

### Study area

The Australian Wet Tropics bioregion is composed of mixed tropical rainforest in an area of approximately 1.85 million hectares. The terrain is rugged and dominated by mountain ranges, tablelands, foothills and a lowland coastal plain. The elevation varies from sea level to highlands at 1000 meters, with isolated peaks reaching up to 1,620 meters [37]. Annual rainfall varies between 1200-8000mm with rainforest covering most areas with annual rainfall above ~1500mm.

### Data collection

Rainforest birds in the Australian Wet Tropics bioregion were monitored between 1996 and 2016 across the region at locations ranging from 0 to 1500 meters above sea level. All individuals were recorded either by call or visual observation. Each survey was based on a 30 minute-150 meter transect with two observers. Surveys occurred within two hours of sunrise. All long-term monitoring sites were located within large, unfragmented areas of rainforest with continuous forest cover over the available elevational gradient. For further details of methods, species observed, site localities and species traits see Williams, VanDerWal (47), Williams, Shoo (50).

Data from four mountain ranges were included in the analyses presented here (Atherton Uplands, Carbine Uplands, Spec Uplands and Windsor Uplands), representing a total of 1977 surveys across 124 different sites. Analysis across these sites was possible based on coverage of the elevational gradient and consistency and numbers of surveys over time. Years with an entire elevation category missing were not included in the trend analyses (from 1996 to 1999). See Table S5 in supplementary information for complete breakdown of survey numbers by elevation and year.

### Species included

Initially, all species for which the survey technique was unsuitable (e.g. water birds) or when call identification was considered unreliable due to the presence of multiple species with similar calls were excluded to ensure maximum reliability in the trend analyses. This resulted in a dataset of 54 species. Of these, analysis of population trends across elevation and time was possible for 42 species where there was sufficient data across both elevation and years to reliably analyse temporal changes in species abundance. We grouped species by habitat (rainforest specialist, generalist) and elevation specialisation (lowland 0-450 m, midland 451-850m, upland >850). These elevational bands were selected to have the finest scale division of elevation possible with a relatively equal band width and sample size in each band. Species were categorised as rainforest specialist if rainforest represented their main or core breeding habitat [47]. Elevation specialisation was defined by the elevational abundance profile of each species using a mean abundance for all surveys conducted between 1996-2004 within each elevational band as the baseline elevation abundance profile (Table S1, Figure S1.1). Species were assigned as a specialist in that elevational band when >70% of their elevational abundance profile occurred within that band.

### Population indices and trends

Overall population indices and trends for all 42 species were modelled using the rtrim package [72], an R-package based on the TRIM software (*Trends and Indices for Monitoring data*. TRIM v. 3.54. [73]). TRIM is designed to analyse time-series of counts and produce unbiased yearly indices and standard errors using log-linear models. The programme also estimates the dispersion factor, correcting for over-dispersion, and takes account of serial correlation between counts at the same site in different years [73]. This method has proved to be robust in trend calculation with missing years [74–77]. We used model 2 in rtrim, which assumed all years as possible changing points in the population trend [78]. Overall trends for each species were calculated with both unweighted data and data weighted by the geographic area within each of the elevational bands. The weighted trends give an estimate of the changes in the species total population size as it takes into account the area within each elevational band as well as local abundance changes. Given that the difference in results using weighted and unweighted data was very marginal (Figure S1) and our primary focus here was to examine site-specific changes in local abundance and elevational distribution shifts in abundance, we have only presented in the results section the unweighted trends. Area-weighted trends of total population size are included in the Appendix (Figure S1b).

Individual species indices produced by TRIM were combined into multi-species indicators for each predefined group. Multi-species indicators were calculated using the MSI-tool in R [79]. This tool uses species-specific annual indices and their standard errors provided by TRIM to calculate annual multispecies indicator with confidence interval, accounting for sampling error, using the Monte Carlo simulation method. This method calculates a mean and a SE from 1000 simulated multi-species indicators and back-transforms these to an index scale, then repeats the process 10000 times. Those indicators are considered a measure of biodiversity change, where a reduction in index mean will occur if more species are declining than increasing and vice versa [75, 80]. We tested for significant differences between the multi-species indicators using the TREND_DIFF-function, based on a Monte Carlo procedure (1000 iterations) and report the average difference with SE in the multiplicative trends and the significance of this difference.

Additionally, we explored the influence of changing trends over the study period by separately examining the trends prior to, and after, 2008 (the midpoint of the time-series). This enables some consideration of how the trends would have been observed over shorter time periods.

Finally, to examine the population changes of each species within each elevational band, we estimated the local trends for each species along the elevational gradient. Multi-species trends in each elevation category were combined for each of the elevation specialist species groups. Differences within groups were tested using the TREND_DIFF-function. R (version 3.6.2) was used in all the analyses [81].

## Acknowledgements

We thank James Cook University, Queensland Department of Environment & Science, Wet Tropics Management Authority, Earthwatch Institute, Rainforest-CRC, Marine & Tropical Research Facility, and the National Environmental Research Program for funding support. We also thank the many volunteers, students and assistants who have contributed to the field work over the years.

